## Supplementary material for "A protease and a lipoprotein jointly modulate the conserved ExoR-ExoS-ChvI signaling pathway critical in *Sinorhizobium meliloti* for symbiosis with legume hosts": SI Figures S1 - S5

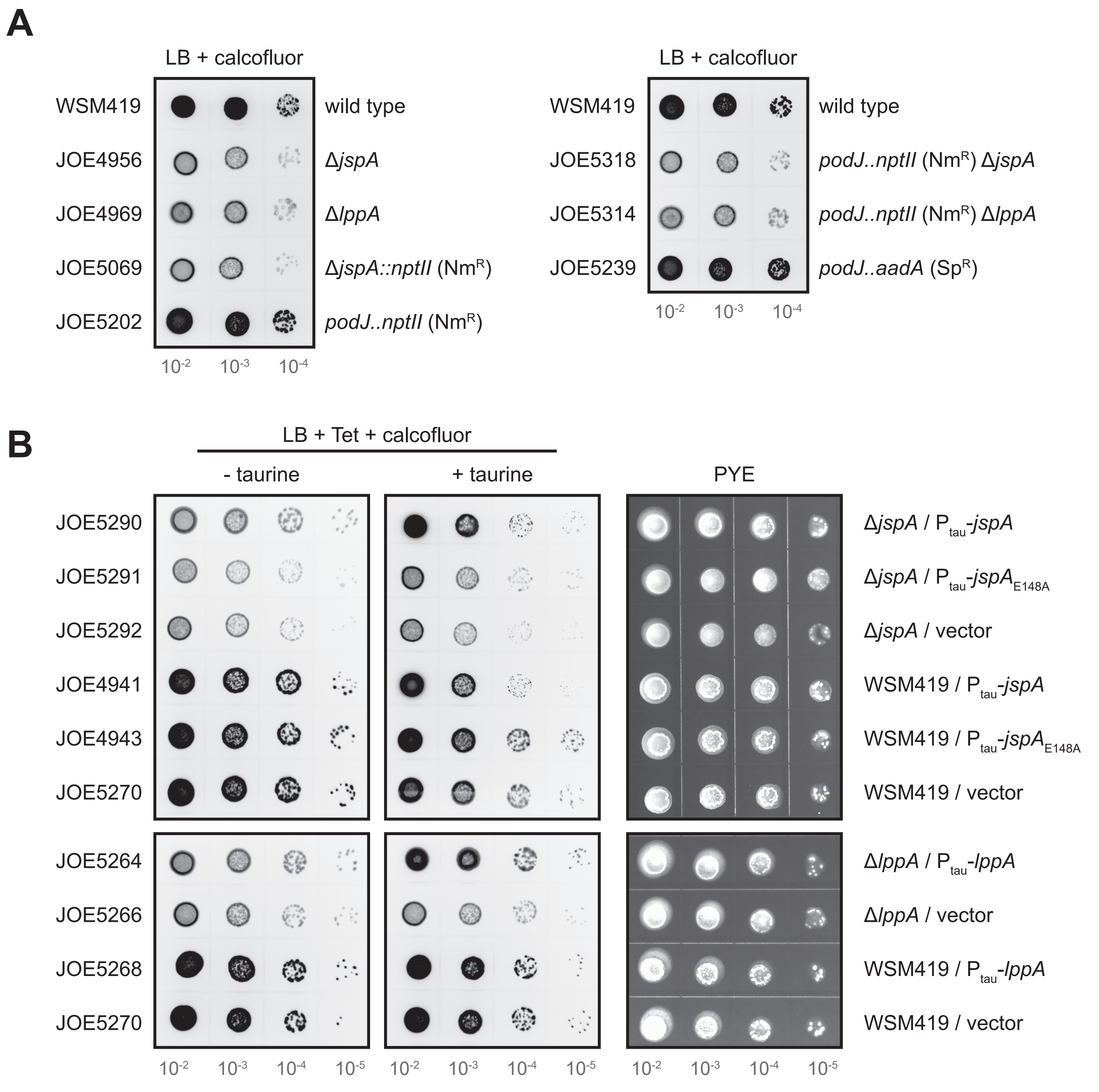


FIGURE S1. Production of calcofluor-binding exopolysaccharides in *S. medicae* WSM419 and its derivatives. Ten-fold serial dilutions of logarithmic-phase cultures were spotted onto solid media and allowed to grow for three days prior to imaging. (A) Representative images show fluorescence of wild-type WSM419, Δ*jspA* (ΔSmed_3110) mutant, Δ*lppA* (ΔSmed_0632) mutant, and derivatives marked with neomycin ($Nm^R$) or spectinomycin ($Sp^R$) resistance [*nptII* or *aadA* linked to *podJ* (Smed_0147) or replacing *jspA*] on LB plates containing calcofluor. Darker spots indicate brighter fluorescence. (B) WSM, Δ*jspA*, and Δ*lppA* strains carrying the vector (pCM130) or a plasmid with *S. meliloti jspA*, $jspA_{E148A}$, or *lppA* under the control of a taurine-inducible promoter (pJC535, pJC555, or pJC532, respectively) were grown on LB plates containing tetracycline (Tet) and calcofluor, without or with taurine (5 mM taurine for *jspA* complementation, 10 mM for *lppA*). Visible-light images of corresponding strains grown on PYE plates show mucoid colonies. Labels on the left indicate strain numbers, while labels on the right indicate genotypes. Each experiment was performed at least twice.

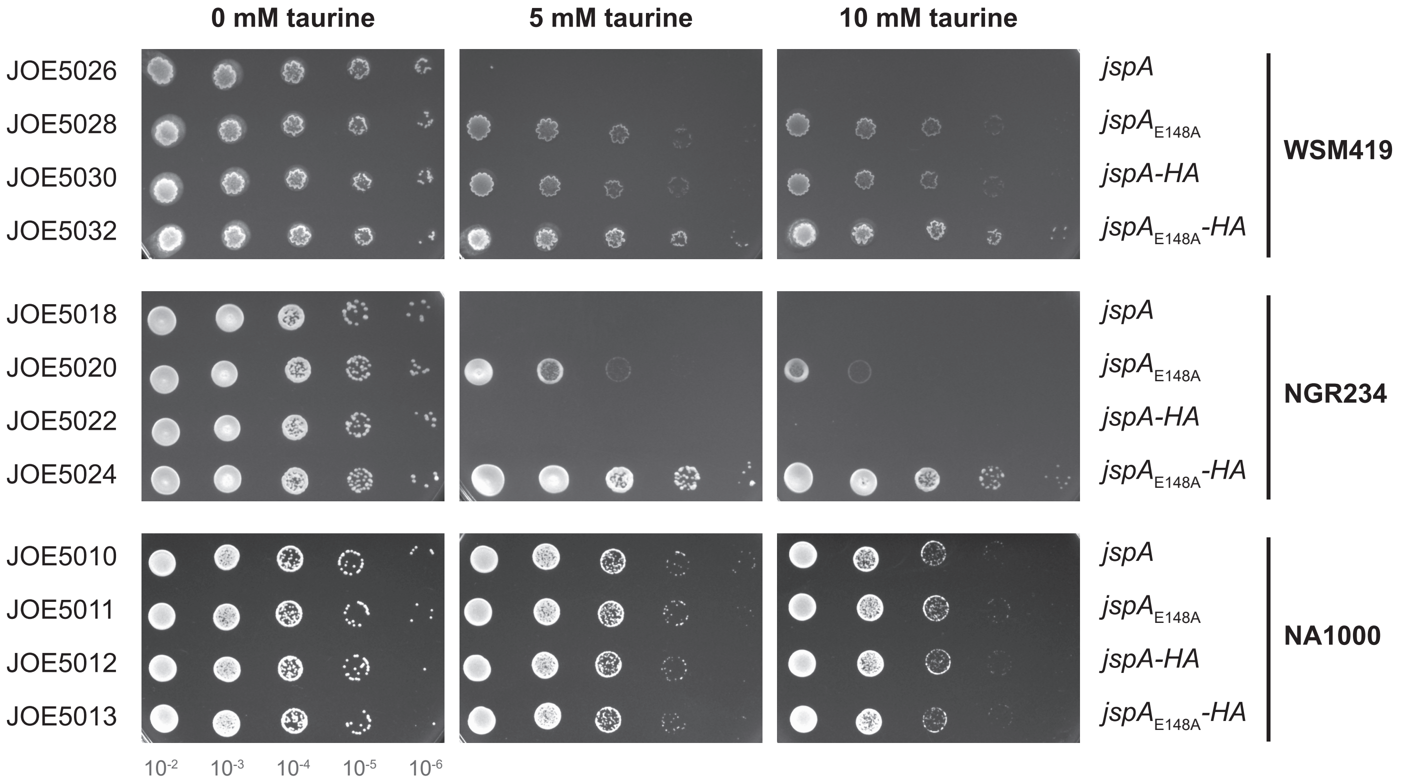


FIGURE S2. Overexpression of *jspA* alleles in *S. medicae* WSM419, *S. fredii* NGR234, and *C. crescentus* NA1000. Ten-fold serial dilutions of logarithmic-phase cultures were spotted onto PYE plates containing 0, 5, or 10 mM taurine. NA1000 strains were grown with 1 µg/mL oxytetracycline for two days, while WSM419 and NGR234 strains were grown with 5 µg/mL oxytetracycline for three days at 30°C prior to imaging. Labels on the left indicate strain numbers, while labels on the right indicate the *jspA* alleles being expressed from a plasmid. Plasmids used were pJC614 (*jspA*), pJC615 (*jspA*$_{E148A}$), pJC616 (*jspA-HA*), and pJC617 (*jspA*$_{E148A}$*-HA*). Images shown represent four replicates on two different days.

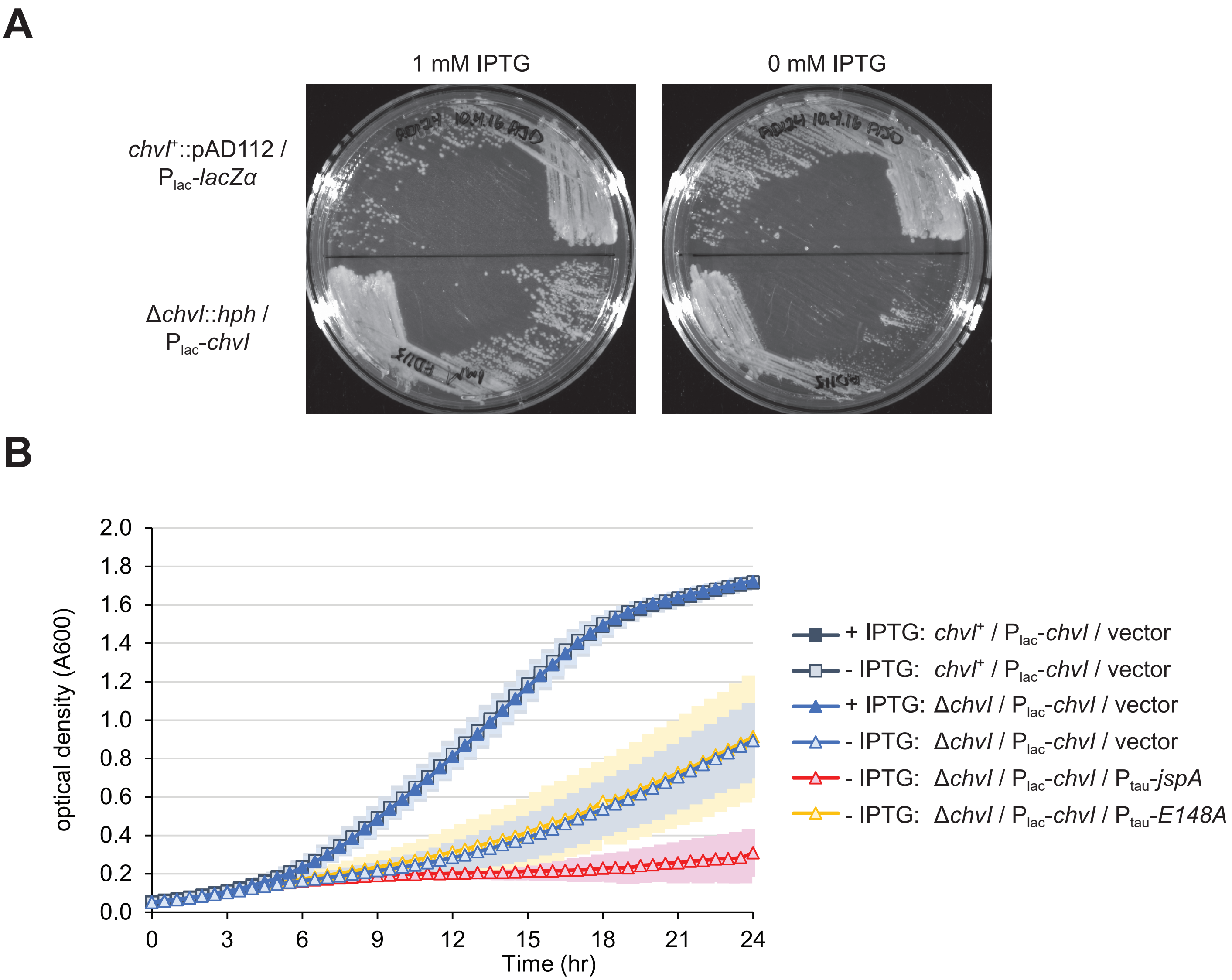


FIGURE S3. Depletion of ChvI. (A) Plate images show growth of ChvI depletion strain on LB medium in the presence and absence of 1 mM IPTG. The top strain (AD124) carried the pSRKKm vector and had a Δ*chvI* allelic replacement plasmid (pAD112) integrated into its chromosome but retained a copy of *chvI*$^{+}$, while the bottom strain (AD115) carried pAD101, with *chvI* under the control of the $P_{lac}$ promoter, and had its *chvI* replaced by a hygromycin resistance gene (*hph*). (B) Plots show growth curves of ChvI depletion strains over 24 hours in LB, in the presence or absence of 0.5 mM IPTG. *chvI*$^{+}$ or Δ*chvI* strains carried pAD101, as well as a compatible vector (pCM130) or derivatives (pJC535 or pJC555) containing taurine-regulated *jspA* or *jspA*$_{E184A}$ (abbreviated as E148A), under control of the $P_{tau}$ promoter. No taurine was added in these growth experiments. Cultures were shaken at 1000 rpm in 48-well plates, with 0.4 mL medium (containing kanamycin and oxytetracycline) per well. Absorbance at 600 nm (A600) was measured every 30 minutes. Average readings for three different days are shown, with corresponding shadings indicating standard deviations. In the presence of IPTG, all strains exhibited similar growth patterns; curves for depletion strains carrying $P_{tau}$-*jspA* or *jspA*$_{E148A}$ grown with IPTG were omitted for clarity. Strains shown here for growth curves (JOE5579, JOE5605, JOE5607, JOE5609) all contain a genomic *exoY-uidA* reporter and constitute a subset of those used for GUS assays in Figure 6C.

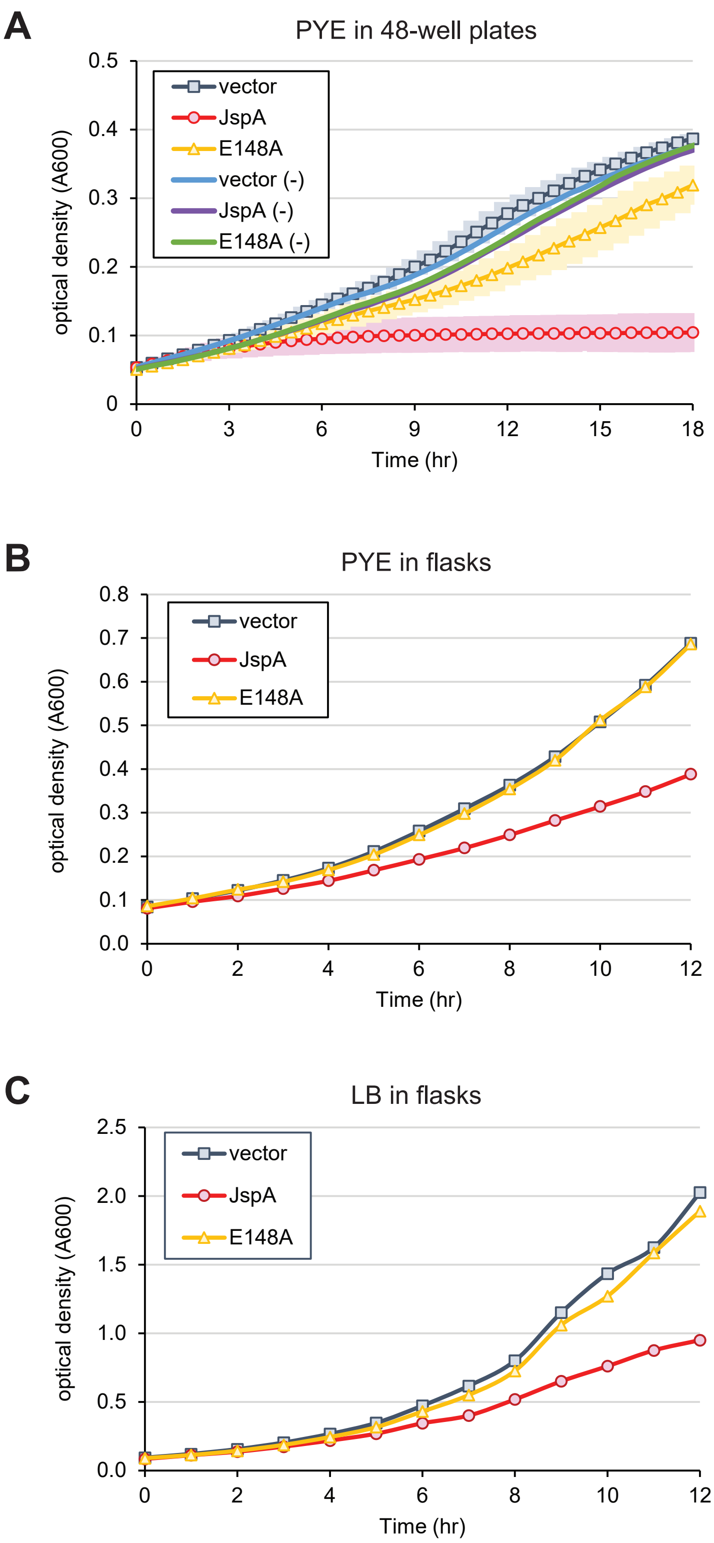


FIGURE S4. Growth curves of *exoR-V5* strains carrying the pSRKGm vector or derivatives (pJC652 or pJC653) with *jspA* or *jspA*$_{E184A}$ (noted as E148A) under control of the $P_{lac}$ promoter. (A) Strains JOE5242, JOE5244, and JOE5246 were grown in 48-well plates, with 0.4 mL PYE per well, in the presence of absence of 1 mM IPTG. Absorbance at 600 nm (A600) was measured every 30 minutes. Average readings for three different days are depicted, with surrounding shadings indicating standard deviations. Lines without markers represent growth in the absence of IPTG (-); standard deviations for these were omitted for simplicity. Figure 7 shows a portion of this graph. (B, C) Liquid cultures of the same strains were grown in flasks with 1 mM IPTG in (B) PYE or (C) LB medium, and A600 was measured every hour for 12 hours. The plots were generated from single experiments.

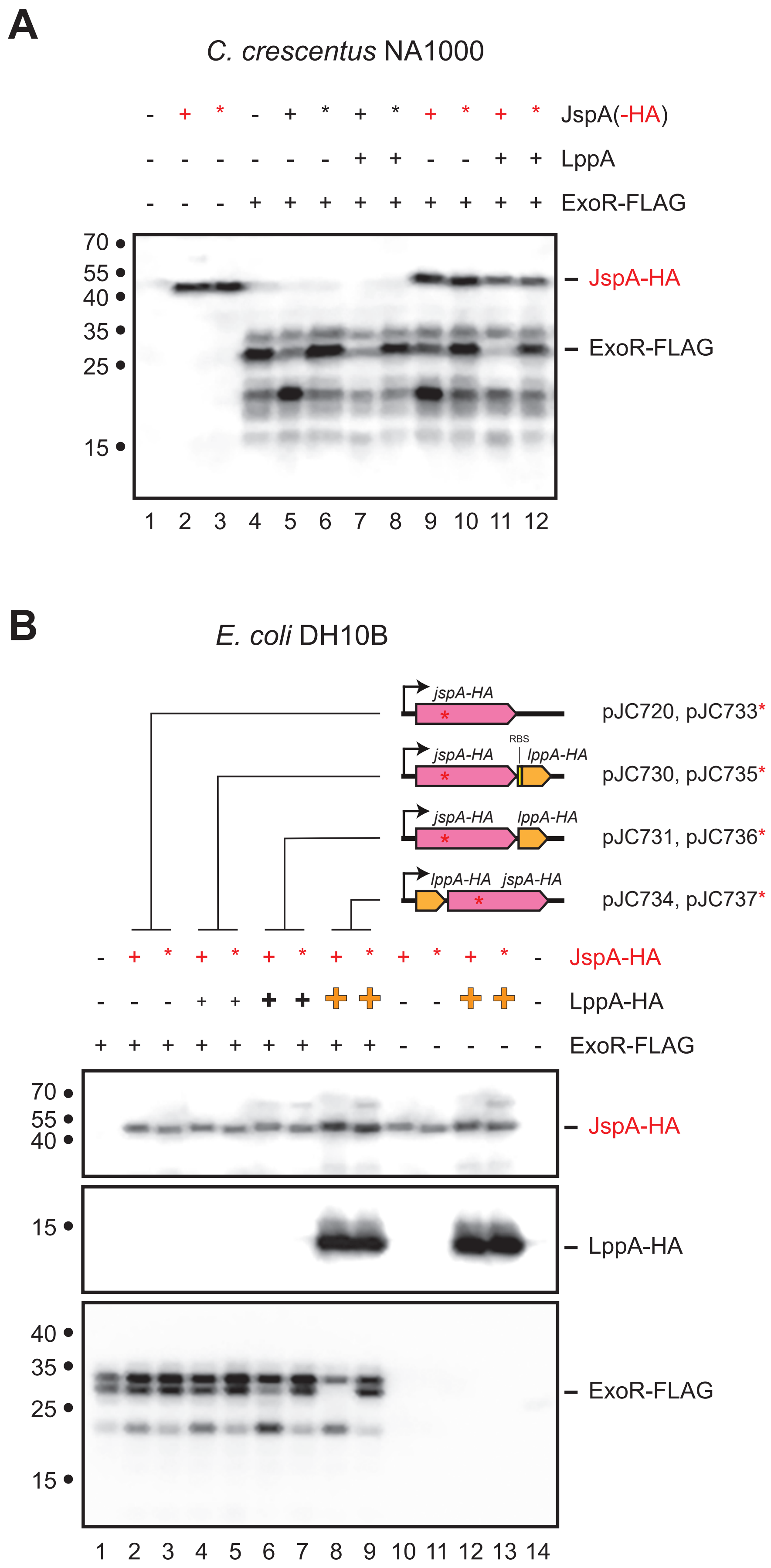


FIGURE S5. Steady-state levels of ExoR-FLAG when co-expressed with different forms of JspA and varying levels of LppA in *C. crescentus* NA1000 or *E. coli* DH10B. (A) Levels of ExoR-FLAG and JspA-HA were assessed in NA1000 by immunoblotting with anti-FLAG and anti-HA antibodies. ExoR-FLAG expression is indicated above the blot: + signifies that expression of ExoR-FLAG from pMB859 was induced with 0.1 mM IPTG in PYE medium for 4 hours, while - signifies that the strain carried the empty vector pSRKKm under the same conditions. Expression of LppA and different versions of JspA were induced with 10 mM taurine from the following plasmids: lanes 1 and 4, empty vector pJC473 when neither expressed (- for both LppA and JspA); lanes 2 and 9, pJC616 for JspA-HA only (red +); lanes 3 and 10, pJC617 for JspAE148A-HA (red *); lane 5, pJC614 for JspA (black +); lane 6, pJC615 for JspAE148A (black *); lane 7, pJC702 for LppA and JspA; lane 8, pJC706 for LppA and JspAE148A; lane 11, pJC707 for LppA and JspA-HA; and lane12, pJC708 for LppA and JspAE148A-HA. (B) Immunoblots show steady-state levels of JspA-HA, LppA-HA, and ExoR-FLAG in DH10B. All strains were induced with 0.1 mM IPTG in LB medium for 4 hours. ExoR-FLAG expression is indicated above the blots: + signifies expression from pMB859, while - signifies carriage of the pSRKKm vector. Expression of different versions of JspA-HA and varying levels of LppA-HA was achieved using different plasmids, similarly indicated above the blots as in (A): lane 1 and 14, pDSW208 vector (expressing GFP); lanes 2 and 10, pJC720 (*jspA-HA* only); lanes 3 and 11, pJC733 (*jspA*$_{E148A}$-*HA*); lane 4, pJC730 (*jspA-HA* and *lppA-HA*, with intervening RBS); lane 5, pJC735 (*jspA*$_{E148A}$-*HA* and *lppA-HA*, with intervening RBS); lane 6, pJC731 (*jspA-HA* translationally coupled to *lppA-HA*); lane 7, pJC736 (*jspA*$_{E148A}$-*HA* translationally coupled to *lppA-HA*); lanes 8 and12, pJC734 (*lppA-HA* followed by *jspA-HA*); and lanes 9 and 13, pJC737 (*lppA-HA* followed by *jspA*$_{E148A}$-*HA*). Schematics above the blots represent gene arrangements on plasmids: red * indicates plasmids that carry the *jspA*$_{E148A}$-*HA* mutant allele and the approximate location of the active site mutation in the gene; RBS preceding *lppA-HA* in pJC730 and pJC735 is the ribosome binding site of *E. coli araB*. Lanes 1 - 7 have the same configuration of strains as those in Figure 9B. The size of the + symbol in the LppA-HA row above the blots reflects the level of expression, with the orange, bold + indicating the highest levels. (LppA-HA is not detectable in some lanes because the signal is overwhelmed by that in lanes with high expression.) Approximate molecular mass, in kDa, are shown to the left of the blots, while lane numbers are shown below. Positions of bands representing JspA-HA, LppA-HA, and ExoR-FLAG are indicated to the right of the blots. Blots were first probed with anti-FLAG antibodies and then with anti-HA antibodies. The ExoR-FLAG image was captured first, and the JspA-HA image was obtained from the same representative blot after the second probing, while the LppA-HA image was acquired from a duplicate blot of the same samples.
