## Supplementary material for "A protease and a lipoprotein jointly modulate the conserved ExoR-ExoS-ChvI signaling pathway critical in *Sinorhizobium meliloti* for symbiosis with legume hosts": SI Tables S1 - S9: JspA SI Table S8 strains.docx

**Table S8.** *Sinorhizobium* strains used in this study

| **Strains** | **Relevant genetic markers, features, and/or description** | **Construction, source, or reference ^a^** |
| --- | --- | --- |
| Rm1021 | SU47 derivative, Sm^R^ (progenitor of strains listed below) | (Meade *et al.*, 1982) |
| Rm7095 | *exoR95*::Tn*5* | (Doherty *et al.*, 1988) |
| Rm7096 | *exoS96*::Tn*5* | (Doherty *et al.*, 1988) |
| CL150 | *ecfR1*^+^ *pstC*^+^ | (Schlüter *et al.*, 2013) |
| CSN015 | Δ*podJ1 lppA*::Tn*5*-110 (SMc00067::Tn*5*-110) | (Fields *et al.*, 2012) |
| CSN134 | Δ*podJ1 jspA*::Tn*5*-110 (SMc03872::Tn*5*-110) | (Fields *et al.*, 2012) |
| EC443 | *exoR-V5* | (Chen *et al.*, 2008) |
| AD084 | CL150 *P_exoY_*-*uidA* (*exoY*::pAD083) | pAD083 mated into CL150 |
| AD113 | CL150 *chvI*::pAD112 (pJQ200sk-Δ*chvI*::*hph*) | pAD112 mated into CL150 |
| AD114 | CL150 *chvI*::pAD112 / pAD101 (pSRKKm-*chvI*) | pAD101 mated into AD113 |
| AD115 | CL150 Δ*chvI*::*hph* / pAD101 (ChvI depletion strain) | Sucrose counter-selection of AD114 |
| AD124 | CL150 *chvI*::pAD112 / pSRKKm | pSRKKm mated into AD113 |
| JOE2612 | *podJ2*(SMc02231)..Ω (Sp^R^ wild-type strain, with antibiotic marker inserted after *podJ2*) | (Fields *et al.*, 2012) |
| JOE3200 | Rm1021 / pCM130 | pCM130 mated into Rm1021 |
| JOE3249 | Δ*jspA* (ΔSMc03872) | (Fields *et al.*, 2012) |
| JOE3366 | Δ*lppA* (ΔSMc00067) | (Fields *et al.*, 2012) |
| JOE3780 | Δ*lppA* *exoS96*::Tn*5* | JOE3366 x Φ(Rm7096), select for Nm^R^ |
| JOE3782 | Δ*jspA* *exoR95*::Tn*5* | JOE3249 x Φ(Rm7096), select for Nm^R^ |
| JOE3902 | Δ*lppA* *exoR95*::Tn*5* | JOE3366 x Φ(Rm7095), select for Nm^R^ |
| JOE3903 | Δ*jspA* *exoR95*::Tn*5* | JOE3249 x Φ(Rm7095), select for Nm^R^ |
| JOE4041 | *lppA*::Tn*5*-110 | Rm1021 x Φ(CSN015), select for Nm^R^ |
| JOE4044 | *jspA*::Tn*5*-110 | Rm1021 x Φ(CSN134), select for Nm^R^ |
| JOE4140 | Rm1021 / pJC535 (pCM130-P_tau_-*jspA*) | pJC535 mated into Rm1021 |
| JOE4400 | Rm1021 / pJC555 (pCM130-P_tau_-*jspA*_E148A_) | pJC555 mated into Rm1021 |
| JOE5242 | EC443 / pSRKGm | pSRKGm mated into EC443 |
| JOE5244 | EC443 / pJC652 (pSRKGm-*jspA*) | pJC652 mated into EC443 |
| JOE5246 | EC443 / pJC653 (pSRKGm-*jspA*_E148A_) | pJC653 mated into EC443 |
| JOE5450 | *P_exoY_*-*uidA* (*exoY*::pAD083) | Rm1021 x Φ(AD084), select for Sp^R^ |
| JOE5526 | *P_exoY_*-*uidA* / pAD101 (pSRKKm-*chvI*) | pAD101 mated into JOE5450 |
| JOE5528 | *P_exoY_*-*uidA* / pSRKKm | pSRKKm mated into JOE5450 |
| JOE5579 | *P_exoY_*-*uidA* / pAD101 / pCM130 | pCM130 mated into JOE5526 |
| JOE5580 | *P_exoY_*-*uidA* / pAD101 / pJC535 | pJC535 mated into JOE5526 |
| JOE5581 | *P_exoY_*-*uidA* / pAD101 / pJC555 | pJC555 mated into JOE5526 |
| JOE5585 | *P_exoY_*-*uidA* / pSRKKm / pCM130 | pCM130 mated into JOE5528 |
| JOE5586 | *P_exoY_*-*uidA* / pSRKKm / pJC535 | pJC535 mated into JOE5528 |
| JOE5605 | Δ*chvI*::*hph P_exoY_*-*uidA* / pAD101 / pCM130 | JOE5579 x Φ(AD115), select for Hy^R^ |
| JOE5607 | Δ*chvI*::*hph P_exoY_*-*uidA* / pAD101 / pJC535 | JOE5580 x Φ(AD115), select for Hy^R^ |
| JOE5609 | Δ*chvI*::*hph P_exoY_*-*uidA* / pAD101 / pJC555 | JOE5581 x Φ(AD115), select for Hy^R^ |
| WSM419 | *S. medicae* isolate from *M. murex* root nodule collected in Sardinia, Italy (progenitor of strains listed below) | (Reeve *et al.*, 2010) |
| JOE4941 | WSM419 / pJC535 (pCM130-P_tau_-*jspA*) | pJC535 mated into WSM419 |
| JOE4943 | WSM419 / pJC555 (pCM130-P_tau_-*jspA*_E148A_) | pJC555 mated into WSM419 |
| JOE4956 | WSM419 Δ*jspA* (ΔSmed_3110) | Allelic replacement using pJC611 |
| JOE4969, 4970 | WSM419 Δ*lppA* (ΔSmed_0632) | Allelic replacement using pJC610 |
| JOE5069 | WSM419 Δ*jspA* (ΔSmed_3110)::*nptII* | Allelic replacement using pJC622 |
| JOE5202 | WSM419 *podJ*(Smed_0147)..*nptII* (Nm^R^ wild-type strain, with antibiotic marker inserted after *podJ*) | Allelic replacement using pJC645 |
| JOE5239 | WSM419 *podJ*(Smed_0147)..*aadA* (Sp^R^ wild-type strain, with antibiotic marker inserted after *podJ*) | Allelic replacement using pJC642 |
| JOE5264 | WSM419 Δ*lppA* / pJC532 (pCM130-P_tau_-*lppA*) | pJC532 mated into JOE4970 |
| JOE5266 | WSM419 Δ*lppA* / pCM130 | pCM130 mated into JOE4970 |
| JOE5266 | WSM419 / pJC532 (pCM130-P_tau_-*lppA*) | pJC532 mated into WSM419 |
| JOE5270 | WSM419 / pCM130 | pCM130 mated into WSM419 |
| JOE5290 | WSM419 Δ*jspA* / pJC535 | pJC535 mated into JOE4956 |
| JOE5291 | WSM419 Δ*jspA* / pJC555 | pJC555 mated into JOE4956 |
| JOE5292 | WSM419 Δ*jspA* / pCM130 | pCM130 mated into JOE4956 |
| JOE5314 | WSM419 *podJ*..*nptII* Δ*lppA* | Allelic replacement in JOE4970 using pJC645 |
| JOE5318 | WSM419 *podJ*..*nptII* Δ*jspA* | Allelic replacement in JOE4956 using pJC645 |

^a^	Φ indicates generalized transduction, as mediated by bacteriophage ΦN3. For example, Rm1021 x Φ(CSN015) means that a bacteriophage lysate made from CSN015 was used to infect Rm1021. ChvI depletion strains were constructed in the presence of 0.5 - 1 mM IPTG.
