## Supplementary material for "A protease and a lipoprotein jointly modulate the conserved ExoR-ExoS-ChvI signaling pathway critical in *Sinorhizobium meliloti* for symbiosis with legume hosts": SI Tables S1 - S9: JspA SI Table S9 plasmids.docx

**Table S9.** Plasmids used in this study

| **Plasmid** | **Relevant genetic markers, features, and/or description** | **Construction, source, or reference ^a^** |
| --- | --- | --- |
| pBAD18 | Arabinose-regulated expression, pBBR322 *ori*, Ap^R^ | (Guzman *et al.*, 1995) |
| pBbB2k-GFP | pBBR1-based broad-host-range plasmid, *gfp* under control of TetR-regulated promoter | (Lee *et al.*, 2011) |
| pCM130 | RK2-derived broad-host-range vector with *E. coli* *rrnB* terminator preceding polylinker, Tc^R^ | (Marx & Lidstrom, 2001) |
| pDSW204 | pTrc99A with weakened P_trc_, *lacI*^q^, Ap^R^, pBR322 *ori* | (Weiss *et al.*, 1999) |
| pDSW208 | pDSW204-*gfp* | (Weiss *et al.*, 1999) |
| pHP45Ω | Source of Ω cassette and *aadA* that confers Sm^R^/Sp^R^ | (Blondelet-Rouault *et al.*, 1997) |
| pJQ200sk | Counter-selectable vector for allelic replacement, *sacB*, Gm^R^ | (Quandt & Hynes, 1993) |
| pSRKGm | pBBR1MCS-derived broad-host-range vector, *lacI*-P_lac_, Gm^R^ | (Khan *et al.*, 2008) |
| pSRKKm | pBBR1MCS-derived broad-host-range vector, *lacI*-P_lac_, Nm^R^/Km^R^ | (Khan *et al.*, 2008) |
| pVO155 | pUC119-derived suicide vector with *uidA*^+^ cassette, Nm^R^/Km^R^ Ap^R^ | (Oke & Long, 1999) |
| pVO205 | pBluescript-derived plasmid, source of *hph* (Hy^R^) cassette | (Barnett *et al.*, 2000) |
| pVO345 | pUC119-derived suicide vector with *uidA*^+^ cassette, Sp^R^ Ap^R^ | (Gibson *et al.*, 2007) |
| pEC340 | pVO155-P*_exoY_* | (Chen *et al.*, 2009) |
| pEC571 | pVO155-P_SMb21188_ | (Chen *et al.*, 2009) |
| pMB694 | pVO345-P*_flaC_* | (Gibson *et al.*, 2007) |
| pMB696 | pVO345 P*_mcpU_* | (Gibson *et al.*, 2007) |
| pMB859 | pSRKKm-*exoR-FLAG* | This study |
| pAD083 | pVO345-P*_exoY_* | This study |
| pAD101 | pSRKKm-*chvI* | This study |
| pAD111 | pJQ200sk-Δ*chvI* | This study |
| pAD112 | pJQ200sk-Δ*chvI*::*hph* | This study |
| pJC382 | pJQ200sk-SMc02231..Ω (for inserting Sp^R^ marker between SMc02231 and SMc02232) | (Fields *et al.*, 2012) |
| pJC440 | pJQ200sk-Δ*jspA* (ΔSMc03872) | (Fields *et al.*, 2012) |
| pJC454 | pJQ200sk-Δ*lppA* (ΔSMc00067) | (Fields *et al.*, 2012) |
| pJC472 | pCM130-P_tau_ | (Mostafavi *et al.*, 2014) |
| pJC473 | pCM130-*tauR*-P_tau_ | (Mostafavi *et al.*, 2014) |
| pJC478 | pCM130-P_tau_-*uidA* | (Mostafavi *et al.*, 2014) |
| pJC532 | pCM130-P_tau_-*lppA* | This study |
| pJC535 | pCM130-P_tau_-*jspA* | This study |
| pJC540 | pVO155-P*_exoR_* | This study |
| pJC555 | pCM130-P_tau_-*jspA*_E148A_ | Megaprimer mutagenesis |
| pJC556 | pCM130-P_tau_-*jspA*_E148D_ | Megaprimer mutagenesis |
| pJC557 | pCM130-P_tau_-*jspA*_H147A_ | Megaprimer mutagenesis |
| pJC558 | pCM130-P_tau_-*jspA-HA* | This study |
| pJC559 | pCM130-P_tau_-*jspA*_E148A_*-HA* | This study |
| pJC560 | pCM130-P_tau_-*jspA*_E148D_*-HA* | This study |
| pJC561 | pCM130-P_tau_-*jspA*_H147A_*-HA* | This study |
| pJC605 | pCM130-P_tau_-*lppA*_C23S_ | Megaprimer mutagenesis |
| pJC606 | pCM130-P_tau_-*lppA-HA* | This study |
| pJC607 | pCM130-P_tau_-*lppA*_C23S_*-HA* | This study |
| pJC608 | pCM130-P_tau_-*lppA*_G96W_*-HA* | PCR error |
| pJC609 | pCM130-P_tau_-*lppA*_A78S_*-HA* | PCR error |
| pJC610 | pJQ200sk-Δ*lppA*_WSM419_ (ΔSmed_0632) | This study |
| pJC611 | pJQ200sk-Δ*jspA*_WSM419_ (ΔSmed_3110) | This study |
| pJC613 | pJQ200sk-Smed_0147 (for inserting markers after *podJ*_WSM419_) | This study |
| pJC614 | pCM130-*tauR*-P_tau_-*jspA* | This study |
| pJC615 | pCM130-*tauR*-P_tau_-*jspA*_E148A_ | This study |
| pJC616 | pCM130-*tauR*-P_tau_-*jspA-HA* | This study |
| pJC617 | pCM130-*tauR*-P_tau_-*jspA*_E148A_*-HA* | This study |
| pJC622 | pJQ200sk-Δ*jspA*_WSM419_::*nptII* | This study |
| pJC638 | pVO155-P*_chvI_* | This study |
| pJC639 | pVO155-P*_pckA_* | This study |
| pJC640 | pVO155-P_SMc01580_ | This study |
| pJC642 | pJQ200sk-Smed_0147..*aadA* (for inserting Sp^R^ marker after *podJ*) | This study |
| pJC645 | pJQ200sk-Smed_0147..*nptII* (for inserting Nm^R^ marker after *podJ*) | This study |
| pJC652 | pSRKGm-*jspA* | This study |
| pJC653 | pSRKGm-*jspA*_E148A_ | This study |
| pJC700 | pCM130-*tauR*-P_tau_-*lppA* | This study |
| pJC702 | pCM130-*tauR*-P_tau_-*lppA-jspA* | This study |
| pJC706 | pCM130-*tauR*-P_tau_-*lppA-jspA*_E148A_ | This study |
| pJC707 | pCM130-*tauR*-P_tau_-*lppA-jspA-HA* | This study |
| pJC708 | pCM130-*tauR*-P_tau_-*lppA-jspA*_E148A_*-HA* | This study |
| pJC715 | pBAD18-*jspA-HA* | This study |
| pJC716 | pBAD18-*jspA*_E148A_*-HA* | This study |
| pJC720 | pDSW204-*jspA-HA* | This study |
| pJC730 | pDSW204-*jspA-HA-*_RBS_*-lppA-HA* | This study |
| pJC731 | pDSW204-*jspA-HA-lppA-HA* | This study |
| pJC733 | pDSW204-*jspA*_E148A_*-HA* | This study |
| pJC734 | pDSW204-*lppA-HA-jspA-HA* | This study |
| pJC735 | pDSW204-*jspA*_E148A_*-HA-*_RBS_*-lppA-HA* | This study |
| pJC736 | pDSW204-*jspA*_E148A_*-HA-lppA-HA* | This study |
| pJC737 | pDSW204-*lppA-HA-jspA*_E148A_*-HA* | This study |

^a^	Plasmid sequence files (in GenBank format) contain construction details. Plasmids pJC608 and pJC609 were generated when missense mutations were introduced during PCR amplification of *lppA* to construct pJC606. Mutant alleles of *jspA* and *lppA* were generated using megaprimer PCR (Tyagi *et al.*, 2004).
